# Differential analysis of genomics count data with edge*

**DOI:** 10.64898/2026.02.16.706223

**Authors:** Lior Pachter

## Abstract

The edgeR Bioconductor package is one of the most widely used tools for differential expression analysis of count-based genomics data. Despite its popularity, the R-only implementation limits its integration with the Python-centric ecosystem that has become dominant in single-cell genomics. We present edgePython, a Python port of edgeR 4.8.2 that extends the framework with a negative binomial–gamma mixed model for multi-subject single-cell analysis and empirical Bayes shrinkage of cell-level dispersion.

## Introduction

The edgeR Bioconductor (Huber et al., 2015) package (Robinson et al., 2010) has been one of the most influential tools in genomics. Together with DESeq2 (Love et al., 2014), it established the negative binomial framework as the standard for count-based differential expression analysis. Since its introduction, edgeR and its underlying statistical ideas have shaped how the field thinks about count data: the negative binomial model for biological variability (Robinson and Smyth, 2007a), TMM normalization (Robinson and Oshlack, 2010), empirical Bayes shrinkage of dispersion estimates (Robinson and Smyth, 2007b), generalized linear models for multifactor experiments (McCarthy et al., 2012), quasilikelihood F-tests (Lun et al., 2016), and gene set testing methods including camera (Wu and Smyth, 2012), roast and fry (Chen et al., 2016). A central theme running through this body of work is the idea, originating in the limma package (Smyth, 2004; Ritchie et al., 2015), that borrowing information across genes via empirical Bayes moderation yields more stable and powerful inference, especially in small-sample settings.

Despite the development of rigorous modeling of technical noise in edgeR, the method was “gene-based”, meaning that the counts analyzed were aggregated over exons or gene bodies. In (Trapnell et al., 2010), the importance of isoform-level analysis was emphasized and a count-based method for isoform differential analysis was introduced in sleuth (Pimentel et al., 2017). The sleuth method relied on estimates of inferential uncertainty, an additional source of noise, to account for the ambiguity of read mapping to isoforms of genes. An adaptation of this approach was incorporated into edgeR in (Baldoni et al., 2024), which led to a method comparable to sleuth and addressed a major flaw of edgeR.

While edgeR now addresses quantification uncertainty, there is another important missing piece that is essential for current genomics applications, namely an approach to single-cell differential analysis. Specifically, single-cell genomics data require assessment not only of technical noise between samples, but also modeling of variability within cell-types. A number of methods have been developed that address this challenge (Finak et al., 2015; Crowell et al., 2020), however such approaches have not been incorporated in edgeR.

To address this shortcoming, we extend the empirical Bayes framework to single-cell data. Multi-subject single-cell differential expression requires a mixed model that accounts for both cell-level overdispersion and subject-level variation. We implement a negative binomial–gamma mixed model following the NEBULA-LN approach (He et al., 2021), and then apply edgeR’s empirical Bayes machinery to shrink the per-gene cell-level dispersion estimates toward an abundance-dependent prior. We implement the extension via the same `squeezeVar` function that underlies edgeR’s quasi-likelihood workflow, but our method is new: neither edgeR nor NEBULA currently perform empirical Bayes moderation of the within-cell dispersion in the mixed model setting.

Rather than working in R, we implement our method in Python, thus providing bidirectional conversion with the popular Python AnnData, which has become a standard for single-cell analysis. AnnData is part of the Scanpy framework (Wolf et al., 2018) and the broader scverse ecosystem (Virshup et al., 2023), and these tools have established AnnData as the standard data structure for single-cell analysis. Most new methods for dimensionality reduction, trajectory inference, and cell type annotation are implemented in Python (Zappia et al., 2018). This creates a practical barrier for researchers who wish to use edgeR’s statistical methods within Python-based workflows: they must either export data to R and import results back, or rely on fragile inter-language bridges that are difficult to maintain and debug. We therefore performed a broad port of edgeR to Python, which we name edgePython, including numerous routines that edgeR relies on written in C^1^.

**Figure 1.**
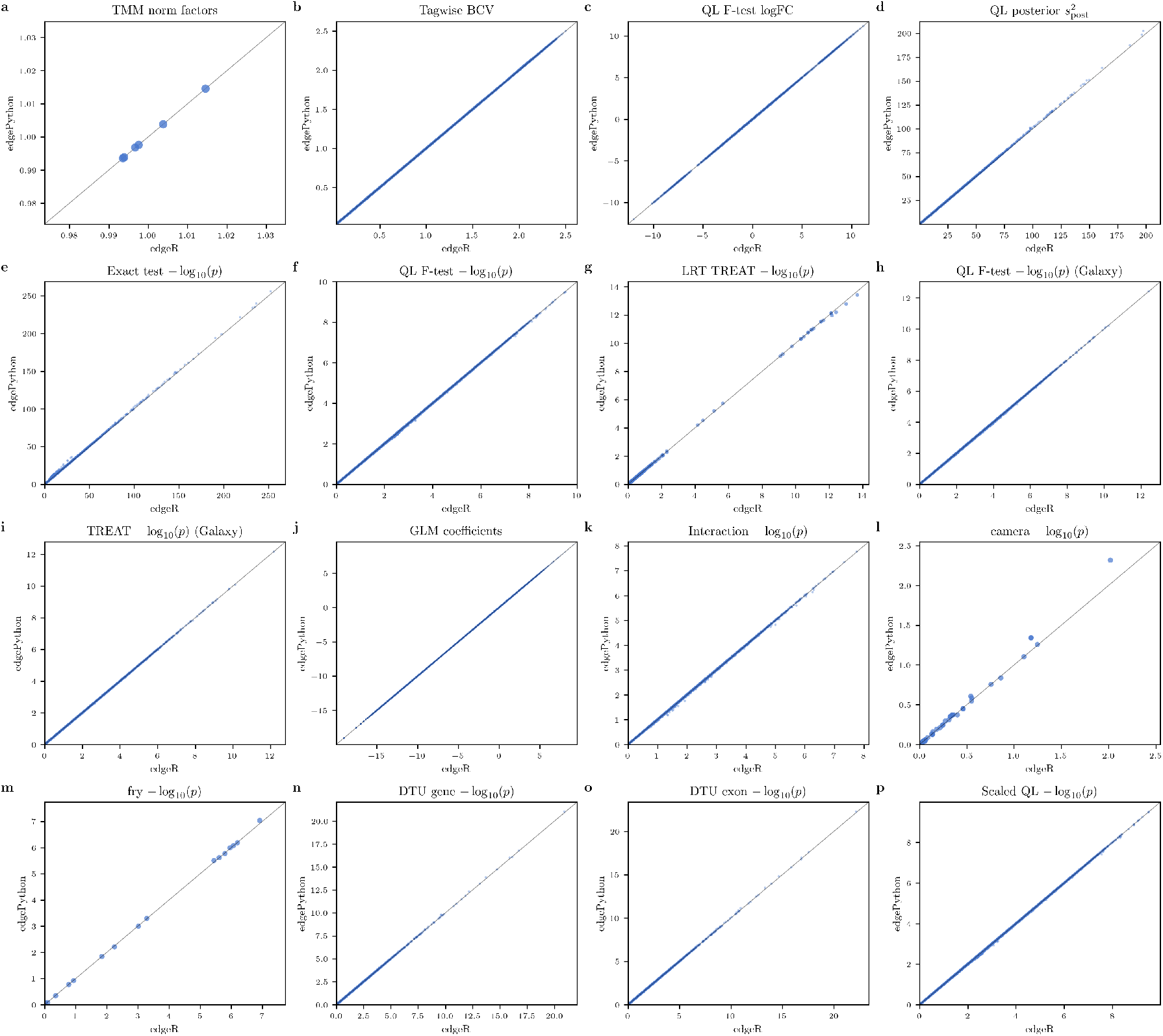
Validation of edgePython. Each panel shows a scatter plot comparing outputs from identical analyses run in R (edgeR, *x*-axis) and Python (edgePython, *y*-axis). Row 1 (a–d): core pipeline components (normalization, dispersion, effect sizes, empirical Bayes shrinkage). Row 2 (e–h): hypothesis testing frameworks (exact test, QL F-test, LRT TREAT, and the GSE60450 multi-factor QL F-test). Row 3 (i–l): multi-factor analyses and gene set testing (TREAT, GLM coefficients, interaction F-test, camera). Row 4 (m–p): additional gene set and specialized analyses (fry, DTU gene-level, DTU exon-level, scaled analysis). **(a)** TMM normalization factors for the HOXA1 dataset (6 samples). **(b)** Tagwise biological coefficient of variation 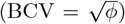 for 44,514 transcripts. **(c)** QL F-test log-fold-changes, validating concordance of effect sizes. **(d)** QL posterior variances 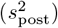 after empirical Bayes shrinkage. **(e)** Exact test *p*-values (−log_10_ scale) for HOXA1 knockdown vs. scrambled. **(f)** QL F-test *p*-values on the same dataset. **(g)** LRT TREAT *p*-values pooled across four log-fold-change thresholds. **(h)** QL F-test *p*-values for the basal pregnant-vs-lactating contrast in the GSE60450 multi-factor design (15,804 genes). **(i)** TREAT *p*-values with log-fold-change threshold 0.58 on the same contrast. **(j)** GLM coefficient estimates (all six coefficients pooled) from the GSE60450 factorial design. **(k)** Multi-degree-of-freedom interaction F-test *p*-values from the same design. **(l)** camera competitive gene set test *p*-values pooled across six configurations. **(m)** fry gene set test *p*-values. **(n)** Differential transcript usage (DTU) gene-level Simes *p*-values for 9,697 genes. **(o)** DTU exon-level *p*-values for 40,958 individual exon tests. **(p)** Scaled analysis QL *p*-values for 44,811 transcripts, incorporating quantification uncertainty via overdispersion scaling. In all panels the diagonal line indicates perfect agreement.

Our port covers all major edgeR functions, including normalization, dispersion estimation, GLM fitting, all four hypothesis testing frameworks (exact test, likelihood ratio test, quasi-likelihood F-test, and treat), as well as gene set testing (camera, fry, roast, mroast, romer). The implementation uses Python dictionaries to mirror edgeR’s S3 list-based objects, and operates on NumPy arrays as well as SciPy sparse matrices. We validate the port by running identical analyses in R and Python and comparing outputs across the full workflow. This should facilitate the incorporation of the edge (empirical analysis of digital expression data) techniques in the computational single-cell genomics ecosystem.

## Results

### Python port

The edgePython program exports 86 public functions and classes organized into 24 modules covering the major workflows of edgeR 4.8.2: data structures (`DGEList, DGEGLM, DGELRT`), normalization (TMM, TMMwsp, RLE, upper quartile), dispersion estimation (common, trended, tagwise, and GLM-based), generalized linear model fitting (negative binomial and quasi-likelihood), all four hypothesis testing frameworks (exact test, likelihood ratio test, quasi-likelihood F-test, and TREAT), gene set testing (camera, fry, roast, mroast, romer, goana, kegga), as well as differential splicing and bisulfite sequencing analysis (Chen et al., 2014). The implementation uses Python dictionaries to mirror edgeR’s S3 list-based objects and operates on NumPy (Harris et al., 2020) arrays and SciPy (Virtanen et al., 2020) sparse matrices. We validated edgePython with 87 unit tests across 15 test files comprising 4,344 lines of test code. These tests cover every major component of edgeR and include direct numerical comparisons against reference outputs generated by running the same analyses in R. Each test verifies that the edgePython matches the edgeR result within a relative tolerance of 10^−3^ or better.

To demonstrate end-to-end concordance on real data, we first applied both edgeR and edgePython to the HOXA1 knockdown dataset (Trapnell et al., 2010) that consists of six RNA-seq libraries (three scrambled controls, three HOXA1 knockdowns) quantified at the transcript level with kallisto (Bray et al., 2016; Melsted et al., 2021; Sullivan et al., 2025), yielding 207,175 transcripts of which 44,514 passed expression filtering. TMM normalization factors computed by both implementations agree to machine precision (1a). Tagwise biological coefficients of variation 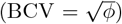 estimated by the full dispersion workflow, including common, trended, and tagwise shrinkage via weighted likelihood empirical Bayes, show near-perfect concordance across all transcripts (1b). The effect sizes also agree: QL F-test log-fold-changes fall on the identity line (1c), and the posterior variances from the empirical Bayes quasi-likelihood 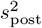 moderation also match closely (1d). The exact test *p*-values for the knockdown effect (1e) and the quasi-likelihood F-test *p*-values from the GLM pipeline (1f) likewise fall on the identity line. Likelihood ratio test TREAT *p*-values, pooled across four log-fold-change thresholds, also agree (1g).

We next applied both tools to the GSE60450 mouse mammary gland dataset, which serves as the basis for a widely used Galaxy RNA-seq differential expression tutorial (Chen et al., 2016). This experiment has a considerably more complex design: 12 samples spanning six groups (basal and luminal cells at the virgin, pregnant, and lactating stages), with a full factorial design matrix. After filtering, 15,804 genes were retained. Quasi-likelihood F-test *p*-values for the basal pregnant-versus-lactating contrast (1h) and TREAT *p*-values with a log-fold-change threshold of 0.58 (1i) show excellent agreement between the two implementations. All six GLM coefficient estimates from the factorial design, including intercept, cell type, two status contrasts, and two interaction terms, match closely when pooled across genes (1j). The multi-degree-of-freedom interaction Ftest, which tests a two-parameter hypothesis, also shows tight concordance (1k). Competitive gene set testing via camera (1l) and self-contained testing via fry (1m) also reproduce across all tested configurations. Two camera points for one gene set deviate modestly; this set contains several genes with very strong differential expression and very low dispersions, making the parametric test statistic sensitive to the same grid-interpolation differences noted above. The rank-based camera variant matches to four decimal places. Finally, we validated two additional analysis modes: differential transcript usage (DTU) at both the gene level (9,697 genes, 1n) and the exon level (40,958 individual tests, 1o), as well as the scaled analysis workflow that incorporates quantification uncertainty via overdispersion scaling for 44,811 transcripts (1p). Across the entire set of comparisons, the maximum observed relative difference in any *p*-value is below 10^−3^.

### Negative binomial mixed model with empirical Bayes shrinkage

Multi-subject single-cell RNA-seq experiments present a statistical challenge not addressed by standard edgeR: the data contain both cell-level overdispersion and subject-level biological variation. Treating individual cells as independent replicates ignores the hierarchical structure and leads to inflated false positive rates (Crowell et al., 2020); a mixed model is needed.

**Figure 2.**
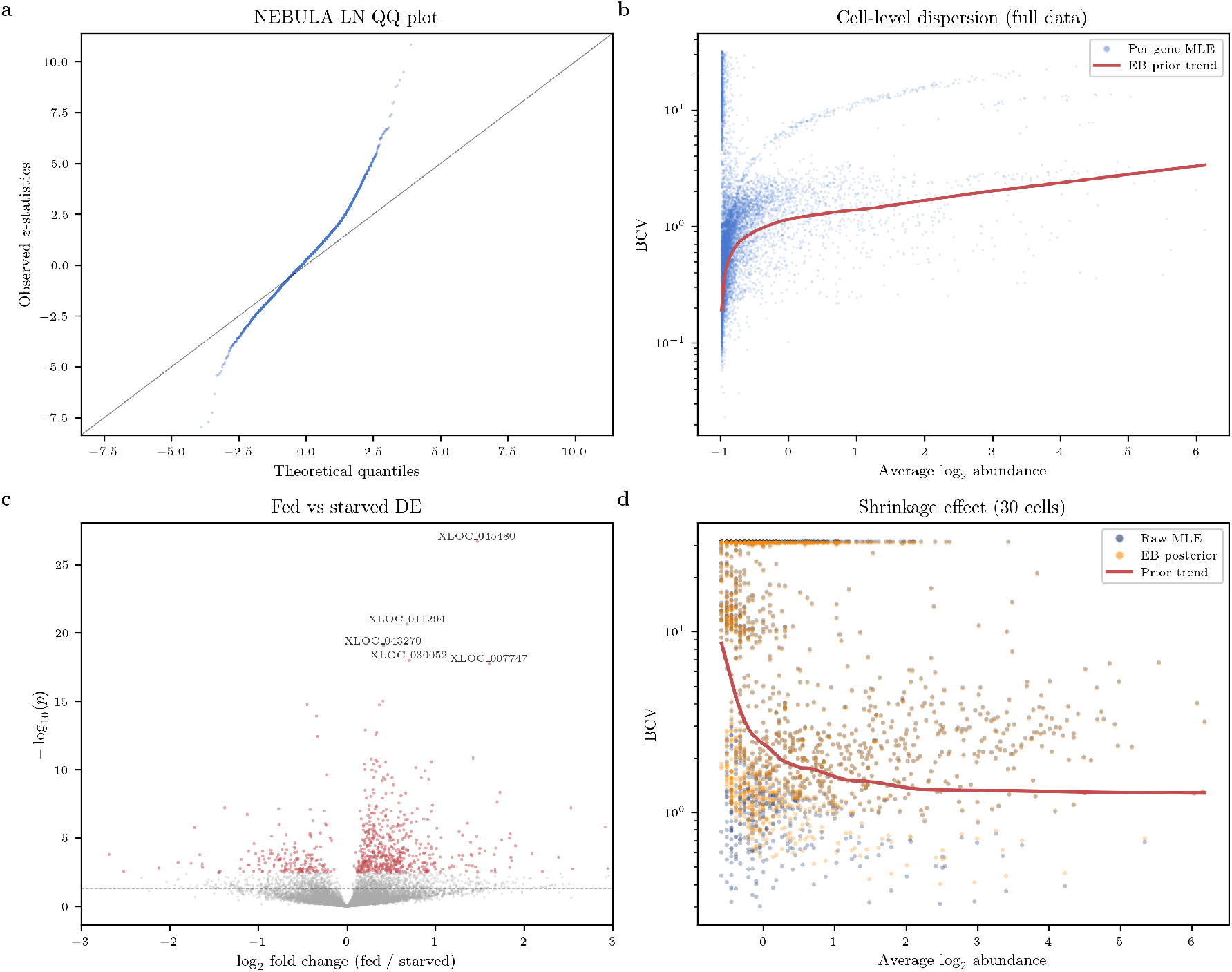
Single-cell negative binomial mixed model with empirical Bayes shrinkage. **(a)** Verification of the NEBULA-LN port: − log_10_(*p*) from R NEBULA (*x*-axis) vs. edgePython (*y*-axis) on the same data. **(b)** Cell-level 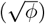 as a function of average log_2_ abundance for 10,377 genes in *Clytia hemisphaerica* gastrodigestive cells (2,564 cells, 10 organisms). The red curve is the empirical Bayes prior trend. **(c)** Volcano plot of fed vs. starved differential expression. Colored points are significant at FDR *<* 0.05; key genes are labeled. **(d)** Effect of empirical Bayes shrinkage on a subsampled dataset (30 cells). Blue: raw MLE dispersions; orange: posterior estimates after shrinkage. The prior trend (red curve) stabilizes the noisy MLEs.

We implement the NEBULA-LN negative binomial– gamma mixed model (He et al., 2021). Let *g* = 1, …, *G* index genes, *j* = 1, …, *J* index subjects, and *i* = 1, …, *n*_*j*_ index cells within subject *j*. The model is

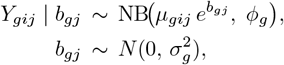

Here, *Y*_*gij*_ is the observed count for gene *g* in cell *i* from subject *j*. The conditional mean is 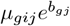, and we use the edgeR/NEBULA negative binomial parameterization

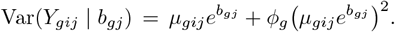

The log-mean model is

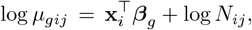

where **x**_*i*_ is the fixed-effect covariate vector for cell *i* (intercept and design terms), ***β***_*g*_ is the corresponding gene-specific coefficient vector, and *N*_*ij*_ is the known library-size offset. The random effect *b*_*gj*_ is a gene-specific subject effect with variance 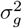, and φ_*g*_ is the gene-specific cell-level overdispersion parameter. TMM normalization (Robinson and Oshlack, 2010) is computed at the pseudobulk (subject) level and expanded to per-cell offsets. Parameters are estimated by a two-stage procedure: L-BFGS-B for initial values followed by penalized maximum likelihood with Newton–Raphson refinement, using a Laplace approximation to integrate out the random effects. The core likelihood and gradient functions are compiled with Numba (Lam et al., 2015) for performance.

The key methodological contribution beyond the NEBULA port is empirical Bayes shrinkage of the cell-level dispersion. The per-gene maximum likelihood estimates 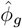 are noisy, especially when the number of cells is modest. We apply the `squeezeVar` function, which is the same function that underlies edgeR’s quasi-likelihood workflow, to borrow strength across genes: the raw 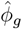 values are shrunk toward an abundance-dependent prior trend, yielding posterior estimates 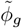 that are more stable. This approach is conceptually identical to the moderation of residual variances in limma (Smyth, 2004), but applied here to cell-level dispersion in the mixed model setting. Neither edgeR nor NEBULA currently performs this shrinkage step.

We verified our NEBULA-LN implementation by comparing the Python output against R’s NEBULA package on the same dataset (2a), confirming that coefficient estimates, standard errors, and *p*-values agree. We then applied the full workflow to *Clytia hemisphaerica* (jellyfish) single-cell RNA-seq data from Chari et al. (2021): 2,564 gastrodigestive cells from 10 organisms (5 fed, 5 starved), with 10,377 genes passing expression filters. 2b shows the per-gene BCV 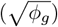 as a function of average log_2_ abundance; the red curve is the empirical Bayes prior trend estimated by `squeezeVar.`

Differential expression analysis between the fed and starved conditions identified 689 genes with significant changes at FDR *<* 0.05 (2c). Of these, 524 are upregulated and 165 are downregulated in fed animals, a 3.2:1 ratio consistent with the expectation that gastrodigestive cells actively upregulate digestive and metabolic gene programs upon feeding. Several of the top differentially expressed genes were not highlighted in the original Chari et al. (2021) analysis, which used a different statistical framework, suggesting that the mixed model approach with empirical Bayes shrinkage can reveal additional biological signal.

To demonstrate the practical value of the shrinkage, we subsampled the data to 30 cells (2d). With fewer cells the per-gene MLE dispersions become markedly noisier, and the empirical Bayes posterior estimates are pulled substantially toward the prior trend. This regularization stabilizes inference in the small-sample regime, where it matters most.

## Discussion

We have presented edgePython, a broad Python port of edgeR 4.8.2 that reproduces the original R implementation across normalization, dispersion estimation, GLM fitting, hypothesis testing, and gene set analysis (1), and extends the framework with a negative binomial–gamma mixed model for multi-subject single-cell differential expression (2). The port of edgeR and its extension are non-trivial. Notably, there has been a previous attempt to port edgeR (edgePy), but the project was ultimately unsuccessful and was aborted with the repository formally archived in January 2025. A project to port DESeq2 (Muzellec et al., 2023) remains active, but the port is not exact, nor does it support all DESeq2 (Love et al., 2014) functions. Although inMoose (Colange et al., 2025) provides a more exact port of edgeR and DESeq2, it is not complete, with three quarters of edgeR’s functionality unavailable despite years of work on the project.

**Figure 3.**
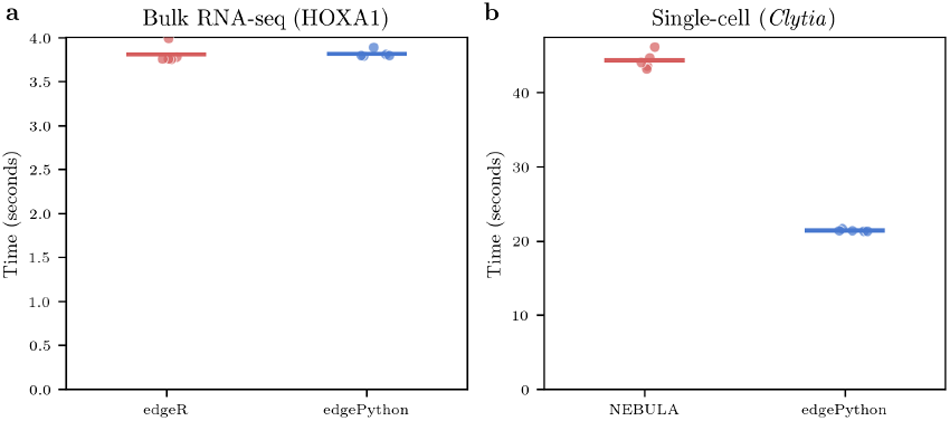
Runtime comparison. Elapsed time for the full differential expression pipeline in R (edgeR/NEBULA) and Python (edgePython) on the HOXA1 bulk dataset and the *Clytia* single-cell dataset.

The performance of edgePython at runtime for bulk RNA-seq analysis is comparable to that of edgeR (3). For the single-cell mixed model, edgePython’s Numba-compiled likelihood and gradient evaluations are substantially faster than R’s NEBULA package. This speed advantage grows with the size of the dataset, making the Python implementation practical for large-scale single-cell atlases.

The edgePython program was developed with the assistance of Claude, a large language model. The faithful translation of a complex statistical package, including numerical optimization routines, special functions, and extensive C code, demonstrates that LLMs can serve as effective tools for cross-language software porting. This suggests that translating edgeR to additional languages is now straightforward: for example, a Rust implementation could integrate with the emerging Rust ecosystem for single-cell genomics or a CUDA implementation could leverage GPUs. The wildcard in “edge*” reflects this idea: the asterisk stands for any target language, given that the barrier to porting has been dramatically reduced by modern language models.

To further reduce the barrier to use, edgePython includes a Model Context Protocol (MCP) server that exposes the full analysis pipeline from data loading through normalization, dispersion estimation, model fitting, hypothesis testing, and visualization. These functions are exposed as tool calls accessible to AI agents, enabling fully automated differential expression analyses driven by natural language instructions. We have verified that agents can successfully run complete bulk and single-cell workflows through this interface.

Our edgePython implementation also provides I/O capabilities beyond those available in edgeR. In particular, kallisto HDF5 output can be imported directly via `catch_kallisto_h5()`, which reads bootstrap samples and estimates per-transcript overdispersion. Although edgeR has supported kallisto bootstrap input via `catchKallisto`, it has not supported HDF5 input, which has been a major barrier for kallisto users. Bidirectional conversion between edgePython’s DGEList and AnnData (`to_anndata()`) or Seurat (`seurat_to_pb()`) objects integrates edgePython into the scverse (Virshup et al., 2023) and Seurat ecosystems (Hao et al., 2024).

Several limitations should be noted: `processAmplicons`, a specialized function for CRISPR screen barcode processing, is not ported. This function reads directly from FASTQ files, and we chose to maintain a clean separation in which edgePython always starts from a count matrix, leaving FASTQ processing to dedicated Python tools. The pathway analysis functions `goana` and `kegga`, which in R depend on organism-specific Bioconductor annotation databases, instead delegate to the g:Profiler REST API (Raudvere et al., 2019), providing a language-agnostic interface with broad organism coverage. The NEBULA-HL (higher-order Laplace) method is not implemented; only the NEBULA-LN approximation is currently available. NEBULA-HL uses second-order corrections to the Laplace approximation to improve the estimation of the subject-level random effects variance, primarily benefiting settings with few subjects or large variance components. However, the empirical Bayes shrinkage of cell-level dispersion that we introduce already stabilizes inference in precisely these small-sample regimes, reducing the marginal benefit of the higher-order correction. On the other hand, the `voom /voomLmFit` workflow (Law et al., 2014), which transforms counts to log-CPM and applies limma-style linear modeling with precision weights, is ported in edgePython. This includes support for within-subject blocking via `duplicateCorrelation` and sample-level precision weights via `arrayWeights`, alongside the negative binomial GLM pipeline.

Finally, it is worth noting a conceptual limitation of count-based differential expression methods in general. edgePython, like edgeR, operates on a single count matrix, yet for single-cell RNA-seq experiments it is now possible and preferable to quantify both spliced and unspliced RNA, yielding two coupled count matrices whose joint distribution is governed by the underlying transcriptional dynamics (Gorin et al., 2023). Methods such as Monod (Gorin et al., 2025) fit biophysically motivated stochastic models to these multimodal counts, enabling inference about transcriptional burst kinetics and regulatory state that a single-matrix framework cannot access. However, there remains a large corpus of legacy datasets for which only a single count matrix is available, and many current assays, including combinatorial indexing protocols, spatial transcriptomics, and CRISPR screens, inherently produce a single layer of counts. For these settings, a well-validated count-based differential expression tool integrated into the Python ecosystem fills a practical need, and edgePython is designed to serve this role.

## Methods

### Software porting

The edgePython program was developed by translating the R source code of edgeR 4.8.2 (Chen et al., 2025) (Bioconductor release 3.20) and its dependencies in the limma, statmod, and locfit packages into Python. The porting was performed with the help of Claude (Anthropic), using the Opus 4.5 and Opus 4.6 models. For each function, the R implementation, including C source when applicable, was provided to the model along with the corresponding R documentation and test cases. Claude produced an initial Python translation that was then validated against the R reference outputs. Iterative debugging resolved numerical discrepancies, typically arising from differences in optimization defaults, special function implementations, or floating-point edge cases. Performance-critical inner loops, particularly in the NEBULA-LN likelihood evaluation and the weighted LOWESS fitting used by the quasi-likelihood pipeline, were compiled with Numba @njit. The complete port comprises 15,318 lines of Python across 24 modules, validated by 87 unit tests (4,344 lines of test code) that compare against R reference outputs.

### Datasets

#### HOXA1 knockdown

Six human RNA-seq libraries (three scrambled controls, three HOXA1 knockdowns) from Trapnell et al. (2010) were quantified at the transcript level with kallisto (Bray et al., 2016; Melsted et al., 2021; Sullivan et al., 2025), yielding 207,175 transcripts. After filtering with `filterByExpr` (default parameters: minimum count 10, minimum total count 15), 44,514 transcripts were retained for analysis. This dataset was used for all panels in 1 labeled “HOXA1” as well as the bulk runtime benchmarks in 3.

#### GSE60450 mouse mammary gland

Twelve mouse mammary gland RNA-seq libraries spanning a 2 *×* 3 factorial design (basal and luminal cells at the virgin, pregnant, and lactating stages; two replicates per group) were obtained from GEO accession GSE60450 (Fu et al., 2015). Gene-level counts for 27,179 genes were filtered with `filterByExpr` to 15,804 genes. This dataset was used for the mouse mammary panels in 1 (panels h–k) and the GLM coefficient comparison (panel j).

#### *Clytia hemisphaerica* single-cell

Single-cell RNA-seq data from Chari et al. (2021) were obtained from CaltechDATA (accession mm6y6-g4569). The complete data set contains 13,673 cells and 28,514 genes. We subset to 2,564 gastrodigestive cells from 10 organisms (5 fed, 5 starved). Genes were filtered by the NEBULA-LN defaults: minimum mean counts per cell *>* 0.005 and expression in ≥ 5 cells, yielding 10,377 genes for analysis.

### Bulk RNA-seq analysis

For the HOXA1 dataset, both edgeR and edgePython were run with identical pipelines: TMM normalization, estimation of common, trended, and tagwise dispersions via `estimateDisp`, and hypothesis testing via exact test, quasi-likelihood F-test, likelihood ratio test with TREAT (at log-fold-change thresholds of 1.0, 1.2, and 1.5), camera, and fry. For the GSE60450 dataset, a full factorial design matrix was fitted using `glmQLFit` with robust empirical Bayes moderation. Contrasts were tested via `glmQLFTest` (single- and multi-degree-of-freedom) and `glmTreat` (log-fold-change threshold 0.58). Differential transcript usage analysis used `diffSpliceDGE` with exon-level F-tests and gene-level Simes aggregation. The scaled analysis incorporated per-transcript overdispersion estimates from kallisto bootstrap samples.

### Single-cell analysis

The NEBULA-LN mixed model was fitted using `glm_sc_fit` with TMM normalization computed at the pseudobulk (subject) level and expanded to per-cell offsets. Parameters were estimated by L-BFGS-B initialization followed by penalized maximum likelihood with Newton–Raphson refinement. Dispersion bounds were set to φ ∈ [10^−4^, 1000] and *σ*^2^ ∈ [10^−4^, 10]. Empirical Bayes shrinkage of cell-level dispersion was performed via `shrink_sc_disp` with robust estimation, using the `squeezeVar` framework with an abundance-dependent prior trend. In 2b, genes whose MLE dispersion hit the optimizer upper bound (φ ≥ 999) were excluded from the plot, as these represent genes too lowly expressed for reliable dispersion estimation.

### Runtime benchmarks

All benchmarks were run on an Apple M1 processor with 16 GB RAM. Each workflow was timed for five-repli cates. The bulk benchmark comprised the full edgeR pipeline (TMM normalization, dispersion estimation, QL F-test) on the HOXA1 dataset. The single-cell benchmark comprised NEBULA-LN fitting on the *Clytia* gastrodigestive subset. R benchmarks used edgeR 4.8.2 and NEBULA 1.5.3. Python benchmarks used edgePython with NumPy 1.22, SciPy 1.11, and Numba 0.58.

### Manuscript

The manuscript was prepared as a PDF/UA-2-compliant PDF using LuaLaTeX with tagged-PDF support via `\DocumentMetadata` and the PDF management/tagging framework in modern TeX Live. Conformance was validated with veraPDF (version 1.28) using the PDF/UA-2 profile.

## Code availability

The edgePython code is available at https://github.com/pachterlab/edgePython. It can be installed with `pip install edgepython.`

## Acknowledgments

The project resulted from a request by Sina Booeshaghi for a Python implementation of edgeR. Thanks to Sina Booeshaghi for also providing invaluable suggestions on how to use Claude effectively and for providing important feedback throughout the project. The work was supported in part by the Caltech Bioinformatics Resource Center.

## Reflection

This project was completed in one week with Claude Opus 4.5 and Opus 4.6 performing the coding. As someone who had not coded seriously for more than 20 years, I found Claude’s efficiency and accuracy enabling in a way that would have been inconceivable a few years ago. At this point, it is reasonable to predict that what took a week for this project may soon take less than a day. This raises the question of what the role of preprints, conference proceedings, and journal publications will be in the scientific enterprise of the future (Zeilberger, 1993).

## Footnotes

1 An analysis of the edgeR code reveals that 39% of the package is written in C, a fraction large enough that edgeR should be called edgeRC.

